# Molecular Identification of *Echinococcus* spp. and other Taeniid Tapeworms Using Next Generation Sequence Analysis of PCR Amplified 18s rRNA gene

**DOI:** 10.1101/2024.04.02.587684

**Authors:** Rasmi Abu-Helu, George Kokaly, Sajeda Nojoum, Imad Matouk, Murad Ibrahim, Ibrahim Abbasi

## Abstract

Cystic echinococcosis (CE) is a prevalent zoonotic disease caused by Echinococcus granulosus, with cosmopolitan distribution. The parasite is transmitted cyclically between canines and numerous intermediate herbivorous livestock animals. Also other taeniid tapeworm could infect domestic dogs and they pose significant veterinary and public health concerns worldwide. This study aimed to develop a sensitive molecular method for detecting Echinococcus spp. DNA in dog fecal samples using next-generation sequencing (NGS). A set of PCR primers targeting conserved regions of Taeniid tapeworms’ 18s rRNA genes was designed and tested for amplifying genomic DNA from various tapeworm species. The PCR system demonstrated high sensitivity, amplifying DNA from all tested tapeworm species, with differences observed in amplified band sizes. The primers were adapted for NGS analysis by adding forward and reverse adapters, enabling sequencing of amplified DNA fragments. Application of the developed PCR system to dog fecal samples collected from Yatta town, Palestine, revealed the presence of *E. granulosus* DNA in five out of 50 samples. NGS analysis confirmed the specificity of the amplified DNA fragments, showing 98-99% similarity with the 18s rDNA gene of *E. granulosus*. This study demonstrates the utility of NGS-based molecular methods for accurate and sensitive detection of Echinococcus spp. in dog fecal samples, providing valuable insights for epidemiological surveillance and control programs of echinococcosis in endemic regions.

**Author Summary:** Cystic echinococcosis, or hydatidosis, is a serious and chronic zoonotic disease in humans caused by the dog tapeworm *Echinococcus granulosus*. The disease is transmitted cyclically between canines and numerous herbivorous livestock animals. Determining *E. granulosus* infection in dogs is crucial for assessing infection risk and identifying new foci of active infections. The infection rate in dogs is also necessary for evaluating transmission dynamics and assessing the efficacy of control programs. In this study, we present a PCR system based on amplification of the 18S rDNA. New primers were designed following an alignment of various taeniid tapeworms’ 18S rDNA sequences. The current PCR system was adapted to be used in amplicon sequencing utilizing next-generation sequencing technology. This strategy enables accurate detection of tapeworm DNA extracted from dogs’ fecal samples and provides quantitative measurement of taeniid infection in dogs.

## Introduction

Cystic echinococcosis (CE) as one of the world’s major zoonotic diseases, which is characterized by the development of large cysts in liver, lungs and other organs of both humans and livestock. The infection is caused by the dog tapeworm, *E. granulosus*, in humans and animals alike (1). Echinococcosis, is one of the most important zoonotic parasitic diseases worldwide and in the Mediterranean region; largely due to extensive home slaughtering practices and presence of numerous stray dogs with access to offal (2-4). The disease is highly endemic in the whole of the Mediterranean Basin and adjacent countries, including Greece, Turkey, Syria, Palestine, Lebanon, Jordan, Egypt, Libya, Morocco, Tunisia and Algeria (5, 6). Reports indicate up to 40% sheep and 30% dogs in Lebanon, and 60% of aged sheep and 20% of dogs in Jordan (7), have been infected. In Palestine, studies on *E. granulosus* are lacking. However, surgical records from West Bank hospitals between 1990 and 1997 a total of 390 revealed 390 surgically confirmed CE cases, with an overall mean annual surgical incidence was 3.1 per 100,000. The highest annual surgical incidence (16.8 per 100,000) was obserev in Yata town near Hebron (8). A study in 2002 found a sero-prevalence of CE among school children in Palestine to be 2.4% (9). Between 2010 and 2015, a total of 353 CE patients were identified were identified from records of 30 hospitals in the West Bank and Gaza (10).

To enable specific identification of *Echinococcus* species, various molecular characterization techniques have been adopted including PCR-RFLP (11) and loop-mediated isothermal amplification (LAMP) (12, 13). However, determining the infection rate in dogs is crucial for epidemiologic studies and surveillance, and control programs, as well as assessing transmission dynamics and infection risks. Traditionally, dog infection has been determined post mortem by identifying worms in intestinal washes or following Arecoline purgation. More recently, an enzyme immunoassay-based coproantigen test has been developed for this purpose, offering genus specific detection with approximately 97% specificity (when worm burdens are more than 50−100 worms) (14, 15). However, this method’s sensitivity for natural *Echinococcus granulosus* infection averages only about 60% (16, 17). To address these limitations and improve detection sensitivity and species-specificity, molecular tools have been developed. These tools have facilitated the differentiation of Taeniidae worms based on morphological and molecular characteristics (18), for identifying *Echinococcus granulosus* and *E. multilocularis* in infected dogs (13, 19, 20). Data accumulation of tapeworms DNA sequences, including mitochondrial DNA, has enabled to develop of molecular diagnostic tools for distinguishing between several human Taenia species (21-24).

Over recent decades, molecular studies, primarily based on mitochondrial genes, have described several genotypes or species within *E. granulosus*. These include *E. granulosus* (Genotypes G1–G3), *E. equinus* (G4), *E. ortleppi* (G5), *E. canadensis* (G6–G10) and E. felidis (‘lion strain’). The existence of the human-specifc genotype G9 is controversial (25). *Echinococcus felidis* had been described in 1937 from African lions, but was later included in *Echinococcus granulosus* as a subspecies or a strain (26) and (11). Further research has clarified the distinction bewteen mitochondrial genotypes, with G1 and G3 now considered as a single species of *E. granulosus*, while G2 is more closely related to G3 (27). Additionally, based on six nuclear loci, G6/G7 and G8/G10 genotypes are considered distinct species (28). Furthermore, it has been proposed that camel and pig strains of *E. granulosus* constitute a single species (*E. intermedius*) (29). Two new species, *E. shiquicus* in small mammals from the Tibetan plateau and *E. felidis* in African lions, , have recently been identified, although their zoonotic transmission potential remains unknown (30).

In our current study we introduced a set of primers based on conserved region found Taeniid tapeworms’ 18s rRNA genes, intended for DNA amplification followed by next generation (NGS) DNA sequencing. The NGS technology allows mass sequencing of genetic material, facilitating the simultaneous production of a vast array of genomic information from multiple organisms amplified by the same PCR reactions in parallel. Moreover, NGS provides a a quantitative counting measurement for each amplified amplicons, reflecting the intensity of infection.

## Material and Methods

### Tapeworms genomic DNA

Previously amplified genomic DNA or newly extracted DNA from various tapeworms from Al-Quds university collection were utilized in this study, encompassing different *Echinococcus* species and *Teania* species.

### Dog fecal samples

A total of 50 samples were collected from household dogs and sheep owner by farmers in Yatta town, situated in the southern part of Hebron district in Palestine. For each collected sample, 5 grams of fecal materials were placed in 50 ml sterile screw-cap tube. Samples were stored at -20°C until DNA extraction.

### DNA extraction

DNA was extracted from dogs’ feces using QIAamp PowerFecal Pro DNA Kit (Qiagen), following manufacturers’ instructions. DNA extraction was performed in duplicate to maximize the concentration of prepared DNA. Approximately 0.2 grams of fecal sample suspended in 100 μl phosphate buffer saline were used each time. DNA was eluted in 50 μl double distilled water and stored at -20°C.

### PCR Primers

A new set of PCR primers was designed based on DNA alignment of tapeworm (*Echinococcus* and *Taenia* species) 18s rDNA genes available in the GenBank. The newly defined primers (Taen18S1D: GGTTTATTGGATCGTACCC and Taen18S1R: CTGTAACAATTATCCAGAGTC) were based on optimally recognized consensus sequences in the aligned regions, spanning a unique sequences for aligned tapeworms 18s rRNA genes, identifiable later by next generation sequencing.

### DNA amplification by polymerase chain reaction

DNA amplification of Taeniid tapeworms’ DNA was conducted using ready to use dry Taq DNA polymerase (Syntezza, Jerusalem). The primers were used in a concentration of 20 pmoles, with approximately 5ng of tapeworms genomic DNA. For PCR amplification using extracted DNA from dogs fecal samples; 5 μl from the extracted DNA was used. The PCR amplification program consisted of an initial denaturation at 95°C for 5 min, followed by 35 cycles (30 seconds at 95°C, annealing at 55°C for 30 seconds, and extension at 72°C for 1 minute). The amplified PCR products were analyzed on 1% agarose gel electrophoresis in TAE buffer (40 mM Tris, 20 mM acetic acid, 1mM EDTA).

### PCR amplification for next generation DNA sequencing

Illumina Miseq (Illumina, USA) NGS strategy, as used for microbiome MiSeq DNA analysis was adopte for DNA sequence analysis (31). This involves two PCR steps, amplification and PCR products index labeling adapted from the bacterial 16s rRNA Metagenomic sequencing library preparation (Illumina, USA). The previously designed primers were modified for NGS analysis adding the following overhang adaptors to the 5’-prime end of each forward and reverse primers: (forward adaptor: TCG TCG GCA GCG TCA GAT GTG TAT AAG AGA CAG, reverse adaptor: GTC TCG TGG GCT CGG AGA TGT GTA TAA GAG ACA G). The PCR products from thew first PCR were individually purified using 0.6X volumes of AMPure XP magnetic beads (AMPure XP beads kit / Beckman coulter, USA), followed by a second PCR reaction to attach the dual indices (S5 and N7) linked to Illumina sequencing adaptors. After index labeling of each PCR product, another DNA amplicon purification was performed using AMPure XP magnetic beads, followed by combining all PCR products into one tube, referred to as the NGS library. NGS DNA sequencing was conducted by an outsourcing service company (sequencing was performed on Miseq machine using 500 cycle kit from Illumina, USA).

### Bioinformatics analysis

Raw Illumina sequencing data were generated from all analyzed PCR amplicons as FASTQ files of read1 (forward) and read2 (reverse) for each individual sample. These sequence reads were uploaded to Galaxy platform at (usegalaxy.org) for further sequence processing and analysis (32). Initially raw sequences were filtered for quality control at a phred score of 20 equivalent to 99% confidence of each nucleotide, followed this forward and reverse reads were merged, the amplified specific genes were selected based on their specific sequence length and sequence identity. The selected sequence reads from each PCR products were analyzed for sequence homology above 97% using BLAST analysis tools.

## Results

### A note on detecting *Echinococcus* parasite’s DNA from dogs fecal samples extracted

The main objective of this study was based was to assess the prevalence of *echinococcus* parasite in dogs using established molecular method (19, 33). Surprisingly, both PCR systems revealed large number of infected dogs upon examination of DNA extracted from collected dog fecal samples. Figure 1 displays the agarose gel electrophoresis analysis of the amplified *E. granulosus* mitochondrial *cox1* gene, clearly indicating 39 positive samples out of the total analyzed 40 samples. The same results were obtained upon retesting the same samples, with an amplification band of 446bp size, using a PCR system targeting a repetitive gene (EgG1 *Hae* III repeat) in the *E. granulosus* genome. As depicted in figure 1, the results of this PCR showed 38 positive samples out of 40 tested. Efforts were made to confirm the positivity of these results through DNA sequence analysis. However, obtaining sequencing information from these bands was challenging, particularly for the second PCR targeting the *E. granulosus* repetitive gene, due to the amplification of multiple bands, rendering conventional Sanger DNA sequencing ineffective. Conversely, the COX1 amplified bands is discrete, but the sequence information was containing many unknown bases (N), indicating several amplicon types with the same band size. Consequently, the search for more specific primers capable of yielding accurate and reliable results without requering further DNA sequence analysis ensued.

**Figure 1:**
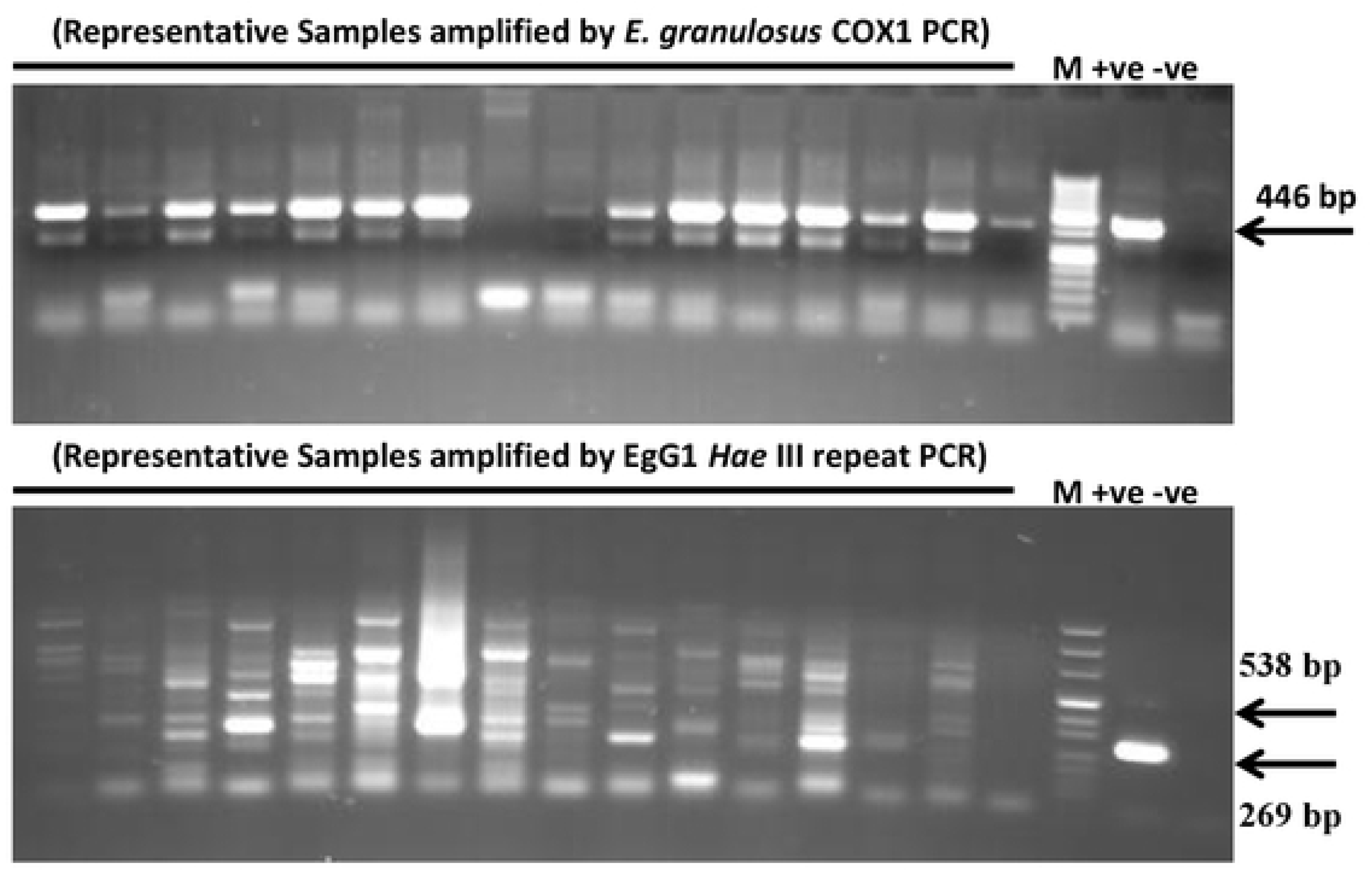
PCR amplification targeting DNA extracted from dogs fecal samples using *E. granulosus* COX1 PCR system (above) and *E. granulosus* (EgG1 *Hae* III repeat) PCR system (lower gel).

### PCR primers used to amplified teaniid tapeworms

Efforts were made to design potential primers for amplifying teaniid tapeworm DNA and conducting sequence analysis by NGS. Different primers were selected based on tapeworm sequences of the mitochondrial and 18s rDNA genes, One set of PCR primers, designed based on 18s rDNA genes demonstrated promise for this purpose. Initially successful in amplifying 5ng of genomic DNA from *E. granulosus, E. multilocularis*, and *Dipylidium caninum* (Figure 2), these primers showed slight differences in the amplified band size, especially when comparing *Echinococcus* species with *D. caninum*. The sensitivity limit of these PCR primers was tested by amplifying pure *E. granulosus* genomic DNA. Demonstrating successful amplification across several twofold dilutions starting from 1 ng/μl, (1, 0.5, 0.25, 0.1, 0.05, and 0.02 ng/μl) with the lowest dilution endpoint not reached (Figure 3).

**Figure 2:**
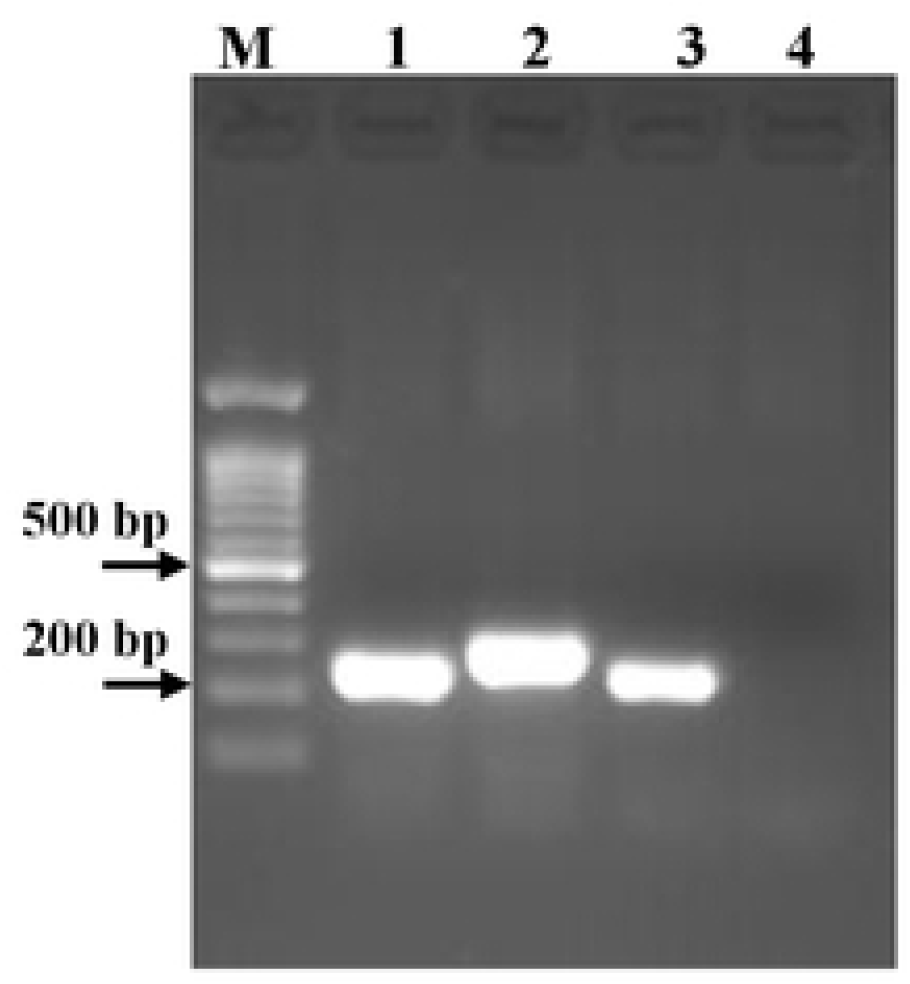
Agarose gel electrophoresis of amplified PCR products from 5ng of genomic DNA. 1-*E. granulosus*, 2- *Dipylidium caninum*, 3-*E. multilocularis*, 4-Negative control. M: DNA size marker.

**Figure 3:**
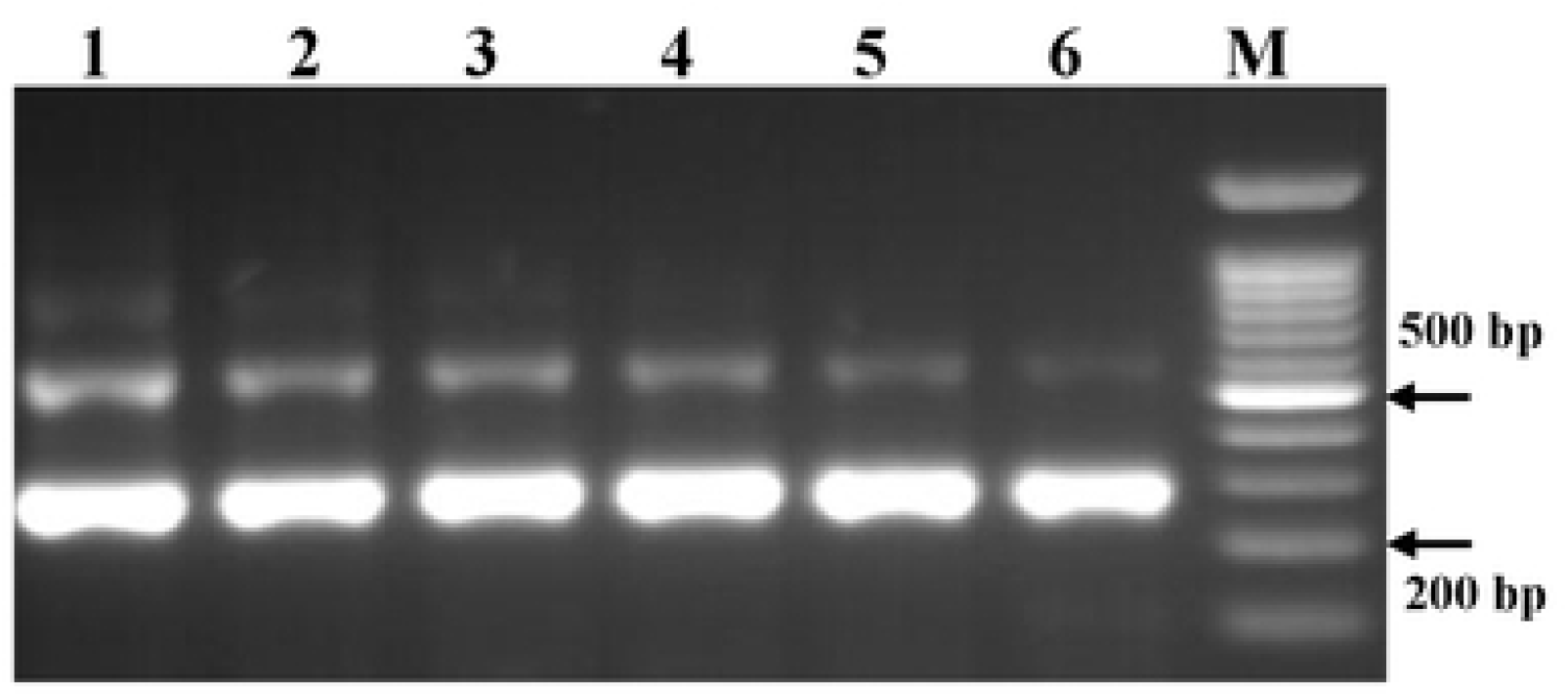
PCR sensitivity test of the three candidate PCR systems. Several dilutions of *E. granulosus* genomic DNA control was used (1, 0.5, 0.25, 0.1, 0.05, and 0.02 ng/ml). M= DNA size marker.

### Adapting the selected primers for tapeworm DNA amplification suitable for NGS analysis

To enable sequencing by NGS methodology, forward and reverse adapters were added to the designed primers. The primers were first used to amplify genomic DNA extracted from various available taeniid tapeworms (*E. granulosus. E. multilocularis, Taenia serialis, T. pisiformis, T. ovis, T. hydatigenia, Multiceps multiceps, D. caninum*). This PCR system that targeting 18s rDNA fragment successfully amplified all the tested species with relatively similar strength. PCR amplification targeting the tested tapeworms and using the designed primers showed slight differences in the amplified bands (Figure 4). Subsequent NGS DNA sequence analysis of the PCR amplified bands precisely identified the exact band size for each of the tested tapeworm species (Table 1). Despite some species exhibiting similar band sizes, NGS sequencing would discriminate between them. Furthermore, the presence of sufficient sequences differences between all tested tapeworm species, with the primers conserved sequences, ensure accurate identification.

**Table 1:** Total number of reads obtained after NGS-DNA sequencing of PCR amplification targeting various tapeworm genomic DNA.

|  |  | Number of reads in tested refernce samples |  |  |  |  |  |  |  |  |  |  |  |  |
| --- | --- | --- | --- | --- | --- | --- | --- | --- | --- | --- | --- | --- | --- | --- |
| Species | PCR amplified band (bp) | S1 | S2 | S3 | S4 | S5 | S6 | S7 | S8 | S9 | S10 | S11 | S12 | S13 |
| *Echinococcus granulosus* | 225/224 | 3266 | 4828 | 0 | 0 | 0 | 0 | 0 | 0 | 0 | 0 | 0 | 1244 | 33 |
| *Echinococcus multilocularis* | 216-218 | 0 | 0 | 5557 | 1679 | 0 | 0 | 0 | 0 | 0 | 0 | 0 | 14 | 11 |
| *Taenia pisiformis* | 263 | 0 | 0 | 0 | 0 | 1684 | 0 | 0 | 0 | 0 | 0 | 0 | 6 | 7 |
| *Taenia multiceps* | 255-259 | 0 | 0 | 0 | 0 | 0 | 2175 | 1277 | 0 | 0 | 0 | 0 | 6 | 31 |
| *Taenia hydatigena* | 267-270 | 0 | 0 | 0 | 0 | 0 | 0 | 0 | 1226 | 0 | 0 | 0 | 67 | 5 |
| *Taenia ovis* | 264-465 | 0 | 0 | 0 | 0 | 0 | 0 | 0 | 0 | 2180 | 0 | 0 | 12 | 19 |
| *Dipylidium caninum* | 230 | 0 | 0 | 0 | 0 | 0 | 0 | 0 | 0 | 0 | 3136 | 2394 | 21 | 2432 |
| Mix of all species (total reads) |  | - | - | - | - | - | - | - | - | - | - |  | 1370 | 2538 |

**Figure 4:**
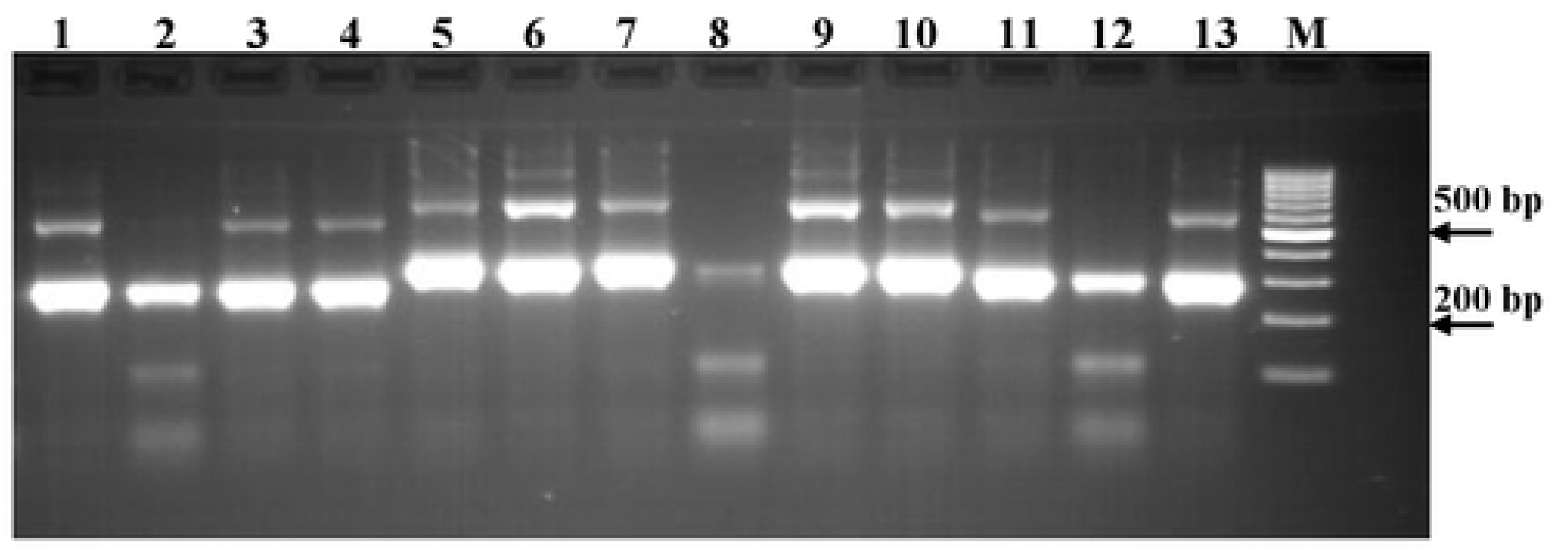
Agarose gel electrophoresis of PCR DNA amplification using the designed primers targeting 1ng genomic DNA extracted from the following tapeworms (1-2: *E. granulosus*. 3-4: *E. multilocularis*, 5: *T. pisiformis*, 6-7: *Multiceps multiceps*, 8: *T. hydatigenia*, 9: *T. ovis*, 10-11: *D. caninum*). M: DNA size marker.

Adapting PCR amplification reaction to next-generation sequence analysis enables quantitative estimation of the amplified PCR amplicons from various tapeworms species. As demostrated in table 1, it was feasible to accurately count the number of amplicons for each of the tested species through NGS sequencing and subsequent bioinformatics counting analysis. The effectiveness of adapting the classically designed primers for NGS sequencing is underscored by the results of PCR amplification of the mixed tapeworm genomic DNA (samples 12 and 13 in table 1).

### Applying PCR systems to DNA extracted from dog-fecal samples

The developed PCR system was applied to amplify DNA extracted from dogs fecal samples. A total of 50 samples were collected from Yatta town were used in this part of the study. Although, PCR results and agrose gel electrophoresis of the amplified products did not allow for calculating the number of positive samples accurately, NGS DNA sequence analysis was imperative for confirmation. The MiSeq NGS was conducted for all the 50 tested samples, generating FASTQ files for each individual sample, including direct and reverse reads files were received (known as Read 1 and Read 2 respectively). These files were processed using Galaxy.org bioinformatics online software, with sequences processed separately for each sample. Based on sequence length and DNA sequence, five samples were found to contain *E. granulosus* DNA amplicons, showing 98-99% similarity with 18s rDNA gene of E. granulosus

## Discussion

*Echinococcus* parasites are tapeworms of significant zoonotic concern, with dogs serving as their definitive hosts, with transmission to humans occurring through ingestion of contaminated food or water. *Echinococcus* tapeworms pose health risks to both dogs and humans, making their accurate identification and effective control imperative for veterinary and public health purposes. Identification of various tapeworm species in dogs’ fecal samples is crucial for effective parasite control and public health management. The most common Taeniid tapeworms found in dogs’ fecal samples include *Taenia pisiformis, Taenia hydatigena*, and *Taenia multiceps, Dipylidium caninum*, with *Echinococcus* species being of particular concern for human health. Dogs become infected after ingesting infected intermediate hosts such as rodents, rabbits, or as in case of *Echinococcus* the ingestion of infected herbivores’ offal. The infective eggs of these tapeworms are shed into the feces of infected dogs. While differentiation between these Taeniid species often relies on the morphological characteristics, molecular techniques such as PCR assays targeting specific DNA sequences have become increasingly valuable for accurate species identification.

Determination of *E*.*granulosus* infection in dogs is crucial for assessing risk of infection, and for identifying active infection foci, and evaluating control program efficacy. Traditionally, identification of *E. granulosus* in dogs has relied on methods such as intestinal washes using arecoline purgation. While coproantigen tests have facilitated large-scale screening of definitive hosts, the development of molecular tools has ben prompted by the need for improved detection sensitivity and species-specific detection. Molecular techniques like PCR not only confirm the presence of the parasite but also provide data on infection intensity and transmission dynamics, enabling effective control measures to mitigate the spread of these zoonotic parasites and predicting disease transmission patterns and mitigating potential public health risks (34, 35).

Molecular diagnosis of Taeniid tapeworms in dogs’ fecal samples has been facilitated by PCR assays targeting specific genetic markers offering high sensitivity and specificity even for low parasite burdens. PCR assays targeting regions of the mitochondrial DNA, such as the cytochrome c oxidase subunit 1 (cox1) gene has been developed for the detection of some Taenia spp (18, 36). Similarly, *Dipylidium caninum* infections haev beed detected through PCR assays targeting the 18S ribosomal RNA gene (37). *Echinococcus granulosus*, identification has been achieved using PCR assays targeting the mitochondrial NADH dehydrogenase subunit 1 (nad1) gene (38). Furthermore, the development of quantitative PCR (qPCR) assays enables the quantification of parasite DNA in fecal samples, offering a potential tool for monitoring treatment efficacy and assessing environmental contamination levels (39).

The 18s rRNA gene, a commonly used for phylogenetic analysis, exhibits a remarkable degree of conservation across diverse taxa, including both Taeniid tapeworms and fungal species. Despite their evolutionary divergence and biological differences, certain regions of the 18s rRNA gene in Taeniid tapeworms and fungal species share notable sequence similarities, reflecting functional constraints on the gene and its role in essential cellular processes conserved across eukaryotes. However, while the overall sequence similarity may exist, specific regions of the 18s rRNA gene can vary significantly, allowing for the discrimination between different taxa. Therefore, while the 18s rRNA gene can be a valuable molecular marker for studying evolutionary relationships and biodiversity, additional genetic markers and complementary analytical techniques are often necessary for accurate taxonomic classification and species identification within these diverse groups.

PCR followed by next-generation sequencing (NGS) offers superior sensitivity and specificity compared to PCR alone for the detection and identification of mixed pathogens in tested samples. PCR amplification enhances the detection of low-abundance sequences, particularly in samples with high background noise or low pathogen concentrations. Subsequently, DNA NGS analysis provides comprehensive sequence information regardless of the mixed amplification of the target DNA, enabling accurate pathogen identification with high sensitivity and specificity (40). This finding is consistent with many studies in the region that estimated dog infectivity by E. granulosus using traditional methods for worm detection, including Arecoline purging tests (41).

In conclusion, incorporating molecular techniques such as PCR followed by amplicon NGS DNA sequencing and bioinformatics analysis into routine diagnostic protocols enhances the accuracy and efficiency of Taeniid tapeworm detection in dogs. These methods not only allow for the differentiation of Taeniid species but also facilitate the detection of co-infections, thereby contributing to improved parasite control and public health management.

